# Regime shifts in a phage-bacterial ecosystem and strategies for its control

**DOI:** 10.1101/797456

**Authors:** Sergei Maslov, Kim Sneppen

**Author notes:** University of Copenhagen, Center for Models of Life, Niels Bohr Institute, 2100 Copenhagen, Denmark. S.M, K.S. contributed equally to this work.

## Abstract

The competition between bacteria often involves both nutrients and phage predators and may give rise to abrupt regime shifts between the alternative stable states characterized by different species compositions. While such transitions have been previously studied in the context of competition for nutrients, the case of phage-induced bistability between competing bacterial species has not been considered yet. Here we demonstrate a possibility of regime shifts in well-mixed phage-bacterial ecosystems. In one of the bistable states the fast-growing bacteria competitively exclude the slow-growing ones by depleting their common nutrient. Conversely, in the second state the slow-growing bacteria with a large burst size generate such a large phage population that the other species cannot survive. This type of bistability can be realized as the competition between a strain of bacteria protected from phage by abortive infection and another strain with partial resistance to phage. It is often desirable to reliably control the state of microbial ecosystems, yet bistability significantly complicates this task. We discuss successes and limitations of one control strategy in which one adds short pulses to populations of individual species. Our study proposes a new type of phage therapy, where introduction of the phage is supplemented by addition of a partially resistant host bacteria.

**IMPORTANCE:** Phage-microbial communities play an important role in human health as well as natural and industrial environments. Here we show that these communities can assume several alternative species compositions separated by abrupt regime shifts. Our model predicts these regime shifts in the competition between bacterial strains protected by two different phage defense mechanisms: abortive infection/CRISPR and partial resistance. The history dependence caused by regime shifts greatly complicates the task of manipulation and control of a community. We propose and study a successful control strategy via short population pulses aimed at inducing the desired regime shifts. In particular, we predict that a fast-growing pathogen could be eliminated by a combination of its phage and a slower-growing susceptible host.

## INTRODUCTION

Diverse ecosystems are known to be capable of regime shifts in which they abruptly and irreversibly switch between two mutually exclusive stable states (1). Such regime shifts have been extensively studied in both macroscopic and microbial ecosystems (1) and shown to be hysteretic and history-dependent. In microbial ecosystems (2) these transitions are known to be possible when a bacterial species directly produces some metabolic waste products or antibiotics (3) that inhibit the growth of other bacteria. They may also occur when bacterial species compete for several food sources, which they use either in different stoichiometric ratios (4) or in different preferential orders (5). Here we explore a new type of regime shifts caused by interactions between bacteria and phages. Bacteriophages have long been known to increase bacterial diversity, especially in aquatic environments (6, 7). However, their potential to create multiple stable states with distinct bacterial species compositions so far has not been recognized. Here we illustrate a possibility of such alternative stable states and regime shifts using a computational model in which two bacterial species compete for the same food source, and are simultaneously exposed to an infection by the same virulent phage. Such dual constraints are known to abate the usual competitive exclusion (8) by allowing multiple bacterial species consuming the same nutrient to co-exist (9, 7).

Microbial communities are an important part of our natural and artificial surroundings and are also responsible for many aspects of human health. Some compositions of microbial communities may be useful for us, while other might be detrimental or even lethal. Thus we would like to reliably manipulate and control the species compositions of these systems. Here we explore several strategies aimed to control the state of phage-bacterial ecosystems via short population pulses inducing the desired regime shift.

## MODEL AND RESULTS

### Model

We study a model describing the dynamics of two microbial species with populations *B*_1_ and *B*_2_ growing on a single limiting nutrient (e.g. carbon source) with concentration *C* and infected by a single phage species with population *P*. All populations are assumed to be well-mixed in an environment constantly supplied with the limiting nutrient at a rate ϕ The dynamics of this ecosystem is given by

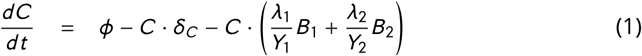

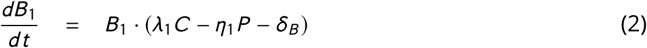

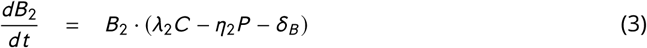

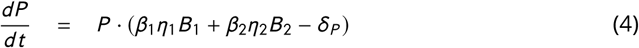

The growth rate of each bacterial species is assumed to be proportional to the nutrient concentration *C* with the species *B*_1_ growing faster than the species *B*_2_: *λ*_1_ > *λ*_2_. Nutrient yields of these two species are given by *Y*_1_ and *Y*_2_ respectively. Phage adsorption coeficients of two species are given by *η*_1_ and *η*_2_ and their burst sizes are *β*_1_ and *β*_2_. The two bacterial species in our model are assumed to have the same death rate *δ*_*B*_ that also includes possible contribution from dilution of their shared environment. The death/dilution rate of the phage is given by *δ*_*P*_ and the nutrient is diluted at a rate *δ*_*C*_.

### Conditions for bistability and regime shifts

In what follows we explore the steady state solutions of Eqs. 1-4, - the only asymptotic dynamical behavior possible in our system. In the absence of phages, the faster growing species *B*_1_ would always eliminate the slower growing species *B*_2_ due to competitive exclusion (8). Phages in principle allow for a slow-growing species to co-exist with the fast-growing one or even to completely take over the ecosystem. In order for this to happen in high-nutrient/high-phage environments the species *B*_2_ needs to be less susceptible to phage infections than the species *B*_1_: *λ*_1_/*η*_1_ < *λ*_2_/*η*_2_. In the extreme case, where the species *B*_2_ is fully resistant to the phage (*η*_2_ = 0), the co-existence between these bacterial species has been previously identified and computationally studied (9, 7, 10).

Here we introduce and study another regime of a phage-bacterial ecosystem in which two bacterial species could mutually exclude each other. This falls under the category of discontinuous and abrupt regime shifts between alternative stable states in microbial ecosystems (see Ref. (2) for a review), which have been previously modelled in the context of competition for nutrients (5, 4) and without phages. In order for a phage-bacterial ecosystem to be in principle capable of bistability, the slow-growing bacterial species needs to produce disproportionately more phages per each unit of consumed nutrient than the fast-growing one: *Y*_2_*β*_2_ > *Y*_1_*β*_1_. As we show in the Supplementary Materials, the bistability requires the following three inequalities to be satisfied:

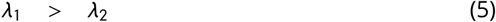

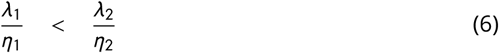

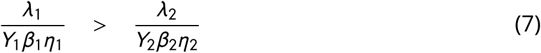

Figure **??**A illustrates the basic mechanisms responsible for bistability and regime shifts in our ecosystem. The thickness of each arrow scales with the relative strength of the interaction between the nodes it connects. Thus the width of the arrow pointing from the nutrient to the bacterial species *B*_*i*_ reflects its growth rate *λ*_*i*_, while that of the arrow pointing in the opposite direction - the rate *λ*_*i*_ /*Y*_*i*_ at which this bacterial species depletes the nutrient. Similarly, the width of the arrow pointing from the phage to the bacterial species *B*_*i*_ reflects its adsorption coefficient *η*_*i*_, while that of the arrow going in the opposite direction - the rate *β*_*i*_ *η*_*i*_ at which this bacterial species generates new phages.

Figure **??**B shows a stochastic simulation of our model with parameters *λ*_1_ = 1, *λ*_2_ = 0.8, *Y*_1_ = *Y*_2_ = 1, *η*_1_ = 0.20, *η*_2_ = 0.15, *β*_1_ = 2, *β*_2_ = 40, *δ*_*C*_ = *δ*_*B*_ = *δ*_*C*_ = 0.2 and *ϕ* = 0.66 (see Methods for details). In our simulations we do not allow the population of either of three species (*B*_1_, *B*_2_, and *P*) to fall below a very small value 4 × 10^−4^. This is equivalent to keeping a constant but weak influx of these species to the ecosystem. As a result, each species would start growing as soon as ecosystem’s internal parameters would make its net growth rate positive.

Random fluctuations in population sizes of bacteria and phages could trigger spontaneous regime shifts between two alternative stable states of the ecosystem visible in Figure **??**B. One of these states is dominated by the fast growing bacterial species *B*_1_. It suppresses the slow-growing species *B*_2_ by the virtue of competitive exclusion via their shared nutrient. In the second stable state the slow-growing species *B*_2_ with a large burst size *β*_2_, generates such a high population of phages that they completely eliminate the fast-growing species *B*_1_, which is relatively more susceptible to phage infections. This steady state also has a larger nutrient concentration due to a slower rate of its depletion by the species *B*_2_.

### History dependence of the ecosystem state

When Eqs. (5-7) are satisfied, the bistability is possible only in a certain intermediate range of the nutrient supply rate. Fig. 2A-D shows the changes in, respectively, steady state values of *P, B*_1_, *B*_2_ and *C* when the nutrient supply rate *ϕ* is slowly changed first up from 0 to 1 and then down to 0 again. For very low nutrient supply rates *ϕ* < 0.04 neither bacteria nor phages can survive and the system stays abiotic *B*_1_ = *B*_2_ = *P* = 0. The fast-growing bacteria *B*_1_ first appears for *ϕ* ≥ 0.04 and prevents the appearance of the slow-growing species due to competitive exclusion. As the nutrient supply rate is increased above 0.14, the population of the phage *P* becomes sustainable and linearly increases with *ϕ*. *B*_2_ continues to be competitively excluded until much higher rate of nutrient supply *ϕ*^(1)^ = 0.70, at which the ecosystem undergoes a regime shift to the state dominated by *B*_2_ and excluding *B*_1_. This alternative stable state persists all the way up the nutrient supply rate. The growth of *B*_1_ is prevented by a high phage population to which this species is especially susceptible. When *ϕ* is lowered, the *B*_2_-dominated state survives down to the nutrient supply rate *ϕ* ^(2)^ = 0.23, which is much lower than *ϕ*^(1)^ = 0.70. Thus for nutrient supply rates between 0.23 and 0.70 the ecosystem is bistable and can be in any of the two alternative stable states making upper and lower parts of the hysteresis loops in Fig. 2A-D. Note that the population of phages and the concentration nutrients generally change in synchrony: when *B*_1_ is dominant, both phage and nutrient levels are low, while the dominance of *B*_2_ generates many phages which significantly lower its population and prevent it from fully exploiting resources, thereby keeping *C* high.

**FIG 1.**
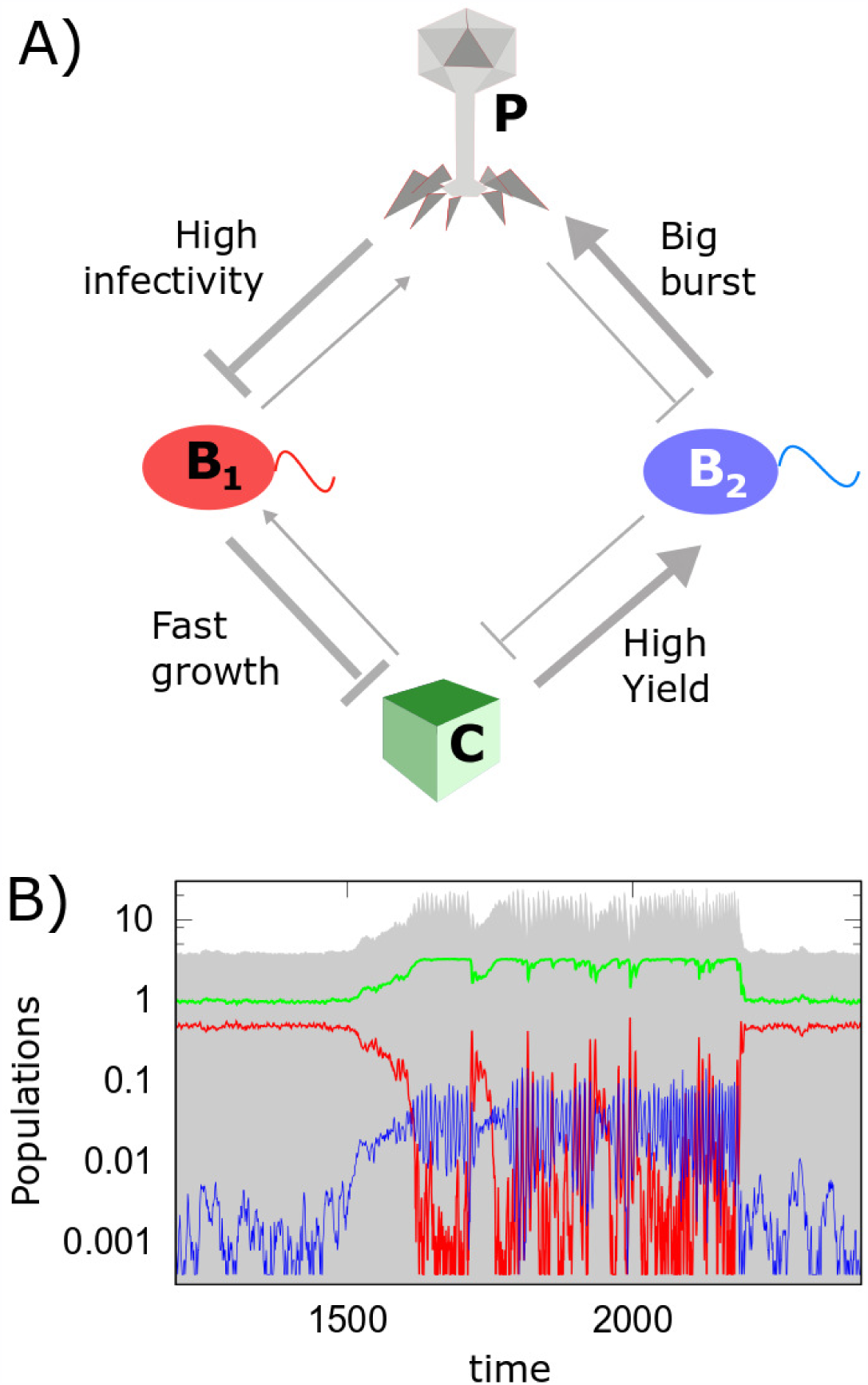
Alternative stable states and regime shifts in a phage-bacterial ecosystem. A) The diagram of interactions between the three species and one nutrient resource in our model: the fast-growing (red, *B*_1_) and the slow-growing (blue, *B*_2_) bacterial species are limited by the same nutrient *C* and infected by the same phage *P*. The slow-growing bacteria are more protected from infections by phage, but, if infected, they generate a larger burst size. The negative effective interaction from *B*_1_ to *B*_2_ is mediated via the nutrient, while that from *B*_2_ to *B*_1_ - via the phage. B) A representative stochastic simulation of the model. Note the abrupt and large regime shifts of the ecosystem between two alternative stable states dominated by bacteria *B*_1_ and *B*_2_ correspondingly. All populations are always maintained above a very low level 4 × 10^−4^ provided by a weak influx of species to the ecosystem. Both phage and nutrient concentrations experience a discontinuous shift up if the ecosystem suddenly flips from the *B*_1_-dominated state to the *B*_2_-dominated one and down in the opposite case. The model parameters are *λ*_1_ = 1, *λ*_2_ = 0.8, *Y*_1_ = *Y*_2_ = 1, *η*_1_ = 0.20, *η*_2_ = 0.15, *β*_1_ = 2, *β*_2_ = 40, *δ*_*C*_ = *δ*_*B*_ = *δ*_*P*_ = 0.2 and *ϕ* = 0.66.

**FIG 2.**
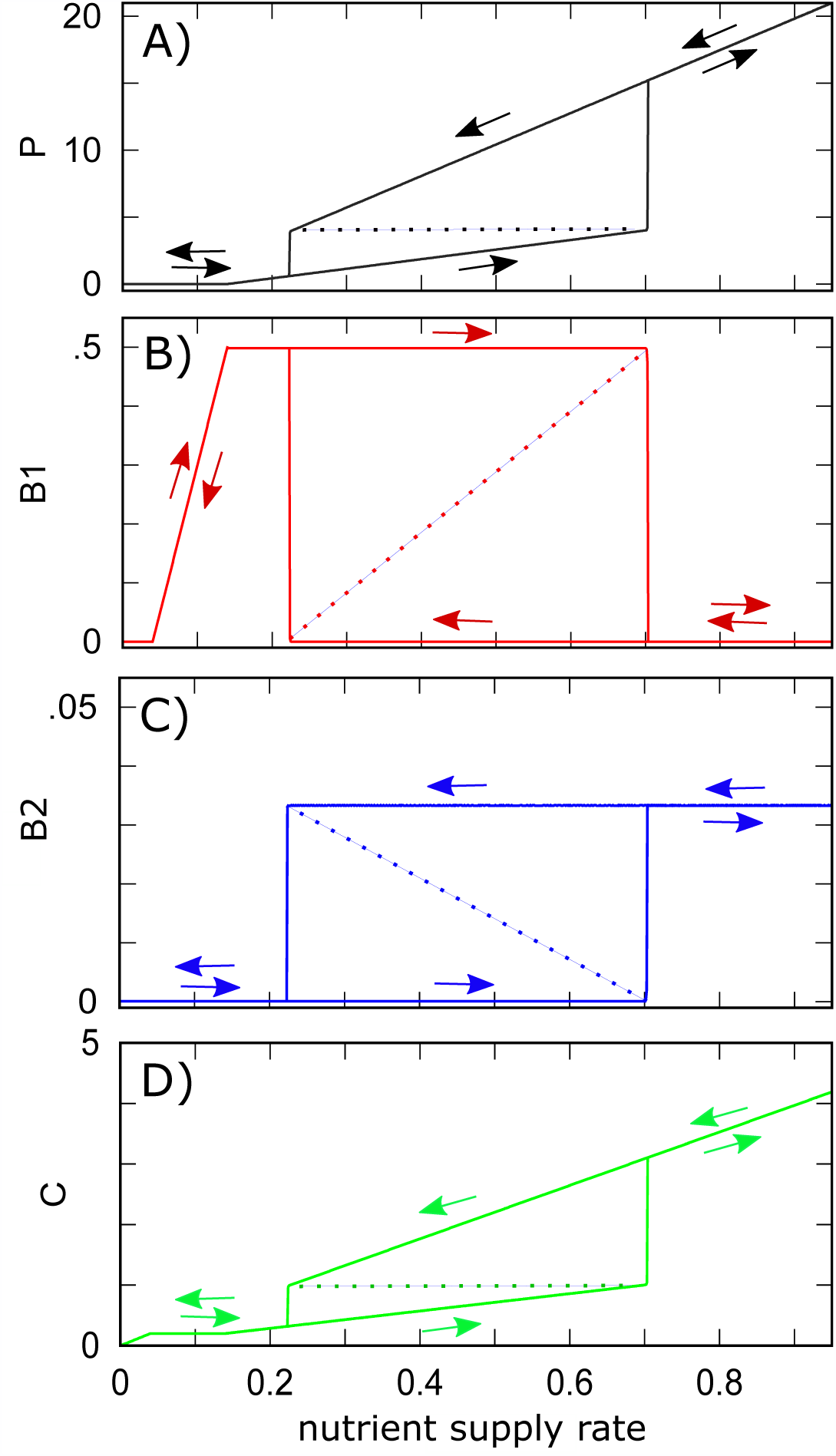
Hysteresis loops in populations of phage *P* (black), fast-growing bacteria *B*_1_ (red), slow-growing bacteria *B*_2_ (blue) and nutrient concentration *C* (green) as the nutrient supply rate *ϕ* (x-axis) is changed first up from 0 to 1 and then down to 0. Note two sudden discontinuous transitions (regime shifts) at both ends of the hysteresis loop. The dashed lines mark populations in the dynamically unstable state separating two alternative stable states. Parameters of the model are the same as in Fig. 1, except for a varying nutrient supply rate *ϕ* (x-axis) and the absence of stochastic fluctuations.

### Controlling regime shifts by population pulses

Phages have recently been investigated as potential agents of control of populations of individual bacterial species in the gut microbiome (11). However, when alternative stable states are present, the state of an ecosystem is complicated by hysteresis and history dependence.

One may need to switch a microbial ecosystem from an undesirable/diseased state to a desirable/healthy state without perturbing the environmental parameters such as nutrient supply rate. One way to achieve such control is by adding a fixed amount of one of the species *P, B*_1_, *B*_2_, or of the nutrient *C* giving rise to an instantaneous increase of its current population/concentration. Such one-time addition, which we call a “population pulse”, is similar to the “impulsive control strategy” discussed in Ref. (12). Since *P, C*, and *B*_2_ are all higher in the *B*_2_-dominated state than in the *B*_2_-dominated state, adding a population pulse of either one of them to the *B*_1_-dominated state could, in principle, trigger a regime shift. Similarly, adding a population pulse of *B*_1_ to the *B*_2_-dominated state could result in a regime shift in the opposite direction.

Fig. 3 explores successes and limitations of the population pulse strategy. We found that this strategy works but only within a certain range of nutrient supply that is generally more narrow than the bistability region itself. A regime shift from the *B*_1_-to the *B*_2_-dominated state can be triggered across the entire bistability region. Conversely, a regime shift from the *B*_2_-to the *B*_1_-dominated state by adding a pulse of *B*_1_ can be made only for *ϕ* below 0.46, which is lower than *ϕ*^(1)^ = 0.7 -the upper bound of the bistable region (the right solid line in Fig. 3). Another observation is the reentrant transition in Fig. 3D: adding too much of *B*_1_ to the *B*_2_-dominated state may prevent the regime shift from taking place. We also note that in order to trigger a regime shift one generally needs to add a pulse that would transiently make the population of the perturbed species to exceed its steady state value in the targeted state (pulse normalized to 1 on the y-axis in Fig. 3). Indeed, a pulse changes only one out of four populations/concentrations in our ecosystem. Thus it needs to be large enough to drive the remaining three populations in the general direction of the regime shift.

**FIG 3.**
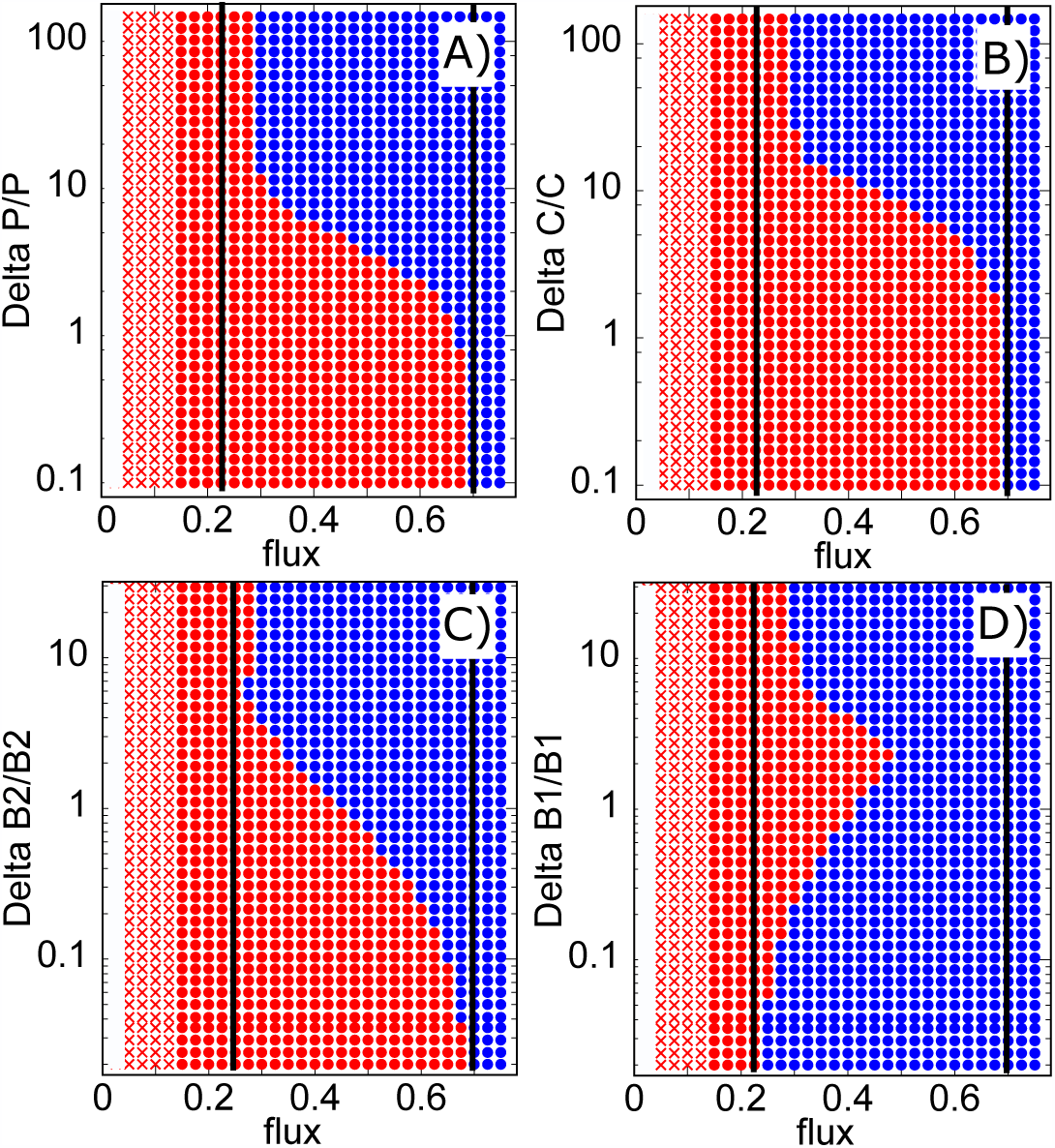
Control of the ecosystem by a pulse in phage population *P* (panel A), resource concentration *C* (panel B), bacterial populations *B*_2_ (panel C), or *B*_1_ (panel D). Red symbols mark the *B*_1_-dominated state, while blue symbols - the *B*_2_-dominated state. In the region marked with red crosses phages cannot exist: *P* = 0. The *x* -axis is the nutrient supply rate *ϕ* with the bi-stable region confined between two black solid lines. The *y* -axis is the magnitude of the pulse normalized by the population/concentration of the target stable state, that is to say, by that of the *B*_2_-dominated state in panels A-C and of the *B*_1_-dominated state in panel D. For nutrient supply rates 0.27 < *ϕ* < 0.7 the *B*_1_-dominated state (red) can be switched to the *B*_2_-dominated state (red) by adding a suffciently large pulse of phage *P* (panel A), nutrient *C* (panel B), or bacteria *B*_2_ (panel C). Conversely, for 0.23 < *ϕ* < 0.46 the *B*_2_-dominated state (blue) can be switched to the *B*_1_-dominated state (red) by adding a sufficiently large pulse of bacteria *B*_1_ (panel D).

Consider a situation where we can simultaneously perturb all three species and the nutrient and set their populations/concentrations (*C, B*_1_, *B*_2_, and *P*) to any desired value. In this case, transient populations after a pulse could be made smaller than their steady state values in the target state. Indeed, to switch the state of the ecosystem, it would be sufficient to make all four populations/concentrations just a little bit closer to the target state than their values in the dynamically unstable state shown as dashed lines in Fig. 2A-D.

### Model with perfect abortive infection in *B*_1_

In one of the phage defense mechanisms called abortive infection (Abi) (13) phages enter and kill the host without producing any phage progeny. A special limit of our model is obtained when the species *B*_1_ is characterized by abortive infection: *β*_1_ = 0, while *η*_1_ > 0. Our equations in this case predict *ϕ*^(1)^ = ∞, which means that *B*_1_ would not disappear from the ecosystem for any nutrient supply *ϕ*. Indeed, this species generates no phage progeny, thus it always can outcompete a small amount of the slower-growing species *B*_2_ infected by phages. However, analogous to Fig. 3A,C a sufficiently large population pulse of *B*_2_ and *P* can get established in the system and eliminate *B*_1_. This could happen for *ϕ* > *ϕ*^(2)^.

## DISCUSSION

We introduced a mathematical model of regime shifts in phage-bacterial ecosystems. The alternative stable states in our model are populated by different bacterial species mutually excluding each other. The negative interactions between these species are mediated by either their co-infecting phages or their shared nutrients. In this respect the mechanism of bistability in our model is similar to that in consumer resource models without phages (4). Indeed, the mandatory (but not sufficient) condition for bistability in either of these two models is a significant difference in stoichiometry of competing microbial species. In our model this stoichiometry is quantified by *Y* · *β* - the product of nutrient yield and burst size of a given bacterial species. It can be interpreted as the conversion factor connecting the amount of nutrients used to build a single bacterial cell to the number phages it produced upon lysis. Comparison of inequalities in Eq. 6 and Eq. 7 shows that bistability is possible only when conversion factors of two bacterial species are sufficiently different from each other: *Y*_2_ · *β*_2_ > *Y*_1_ · *β*_1_.

Similarly, multistability studied in Refs. (14, 4) requires species competing for two types of essential resources (e.g. C and N) to have different C:N stoichiometries.

Regime shifts and multistability are known to occur when competition between species in principle allows for their co-existence, while the differences in stoichiometry make such coexistence dynamically unstable (14, 4). This is also true in our model, where bistability between species *B*_1_ and *B*_2_ is possible whenever their co-existence is dynamically unstable. Conversely, a dynamically stable co-existence of *B*_1_ and *B*_2_ is possible whenever inequalities given by Eqs 5-6 are satisfied, while that in the Eq. 7 changes the direction to *λ*_1_/(*Y*_1_*β*_1_*η*_1_) < *λ*_2_/(*Y*_2_*β*_2_*η*_2_).

Our model predicts that regime shifts in phage-microbial ecosystems can be a consequence of differences in species’ yields *Y*_2_ > *Y*_1_ rather than their burst sizes. A negative correlation between species’ growth rate and its yield known as rate-yield trade-off is widely known (15). According to this correlation slower growing species tend to have higher yields thereby facilitating bistability in our model.

A general case of predator-prey food webs with multiple trophic levels has been considered in Ref. (16, 17). For certain combinations of parameters one can prove that the steady state of dynamical equations describing such ecosystems is unique and thus multistability is impossible. This proof, based on the Lyapunov function proposed in Ref. (18), requires the food web to have identical stoichiometry products (like *Y*_*i*_ *β*_*i*_ in our model) for all paths connecting the same pair of species. Here we extend this study by showing that if the difference in stoichiometries of two such paths is sufficiently large, multistability could in principle emerge. Thus, it is tempting to extend our mechanism for multistability up from microscopic phage-bacterial ecosystems to macroscopic predator-prey food webs. In order for macroscopic food webs to be multistable, the biomass conversion ratio between two successive trophic levels has to deviate widely from its typical value of about 10% (19, 20) and be sufficiently different for different species in the same trophic level. Indeed, one could always choose to measure the population of each species in units of its biomass per unit area. These units would rescale absolute values of competition parameters such as *λ* and *η*. In these units stoichiometric coefficients *Y* and *β* are given by the efficiency (0%-100%) of biomass conversion between two consecutive trophic levels. Multistability requires sufficient differences in biomass conversion factors along paths between species in different trophic levels. For example, in our model the nutrient, which can be thought to occupy the trophic level 0 is connected to the phage species (trophic level 2) via paths going through two different bacterial species (intermediate trophic level 1). Furthermore, the number of species in intermediate trophic levels of these paths has to be odd. Given that the overall number of trophic levels rarely exceeds 4, the case of a single intermediate trophic level considered in this study represents the most biologically plausible scenario.

The ecosystem used in our study is very simple: it has low species diversity and a single growth-limiting nutrient. This simplicity allowed us to quantitatively understand the principal mechanisms giving rise to bistability. More complex ecosystems with a larger number of species and multiple nutrients are expected to have qualitatively similar properties. They also could have a much more complicated phase diagram in the space of nutrient supply rates. Hence multistability with more than two stable states could be realized in some regions of this space (See Ref. (4) for this type of multistability in consumer resource models). Another limitation of our model is that it ignores the possibility of rapid evolution of bacterial strains competing with phages. Such red queen dynamics often generates phage-resistant bacterial strains. The appearance of a phage-resistant variant of *B*_1_ would modify the behavior of our ecosystem for very high nutrient supply, but might not affect bistability between *B*_1_ and *B*_2_ for intermediate nutrient supply studied above. This depends on the magnitude of the growth deficiency of the resistant mutant. A delicate interplay between multiple strains and species could be understood by visualizing them all in Fig. 4A, where a phage-resistant strain would be shown as a vertical line.

**FIG 4.**
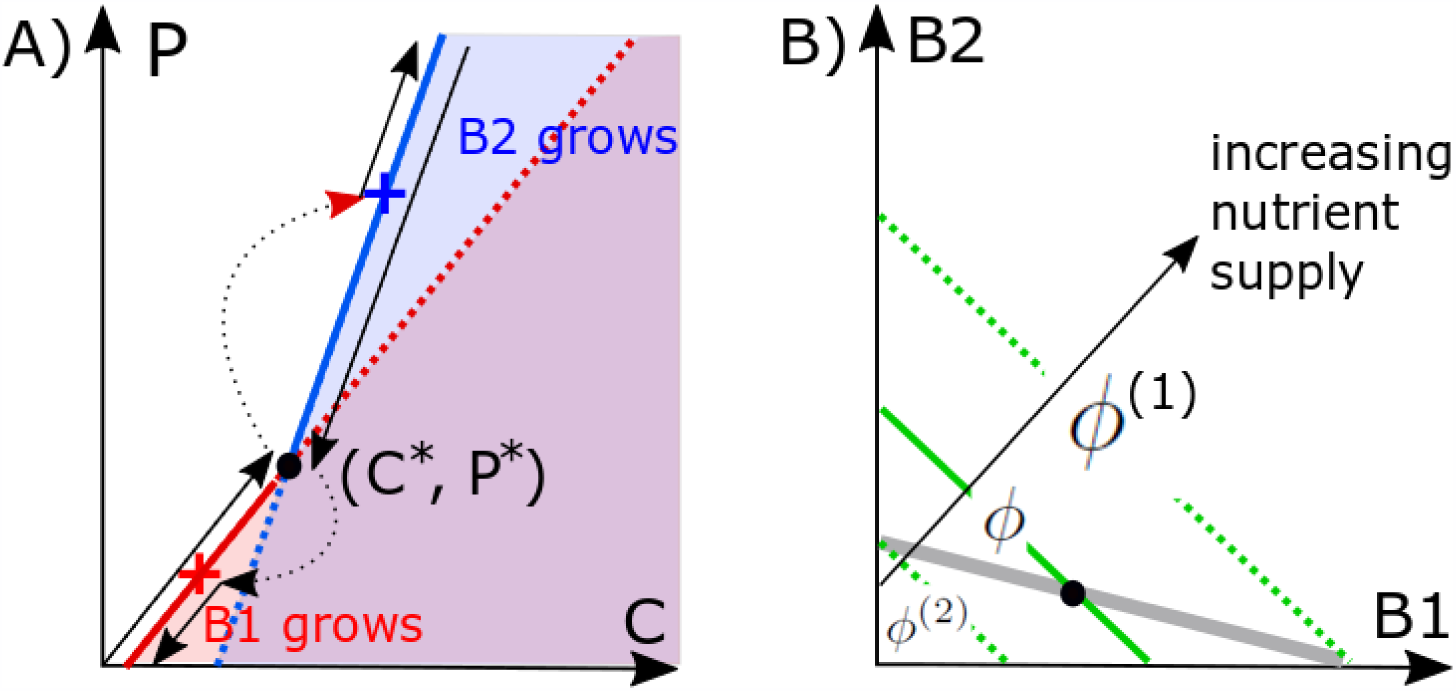
Geometric solution of the steady state of the ecosystem. A) Steady state *C* and *P* are from solving Eqs. (2-3). When both bacteria *B*_1_ and *B*_2_ are present the system can only be at the intersection (*C* ^*^, *P* *). In our case this state is dynamically unstable. As *ϕ* increases, the environmental parameters (*C, P*) follow the solid red line up to the black dot, then discontinuously jumps to the blue cross and continues up along the solid blue line. When *ϕ* subsequently decreases in the hysteresis loop shown in Fig. 2, the (*C, P*) follow the solid blue line down to the black dot, discontinuously jumps to the red cross and continues down along the solid red line. This trajectory is shown in black lines with arrows. B) The geometric solution for coexisting bacterial populations is given by the intersection of the grey line, where the phage population is at the steady state *P* = *P* ^*^ (Eq. 14), and the green line, where the nutrient concentration is at the steady state *C* = *C* ^*^ (Eq. 15). The green line shifts up as the nutrient supply *ϕ* is increased. Bacterial populations *B*_1_ (*B*_2_) disappear at the boundaries *ϕ*^(1)^ (*ϕ*^(2)^) of the bistability region *ϕ*^(2)^ < *ϕ* < *ϕ*^(1)^. Here we show an example in which the steady state *C* ^*^, *P* ^*^ is dynamically unstable giving rise to bistability. However, if the grey line has a steeper slope than the green line, the bistability is replaced by the region (*ϕ*^(1)^ < *ϕ* < *ϕ*^(2)^) of stable coexistence of *B*_1_ and *B*_2_.

It is instructive to compare the mechanisms of bistability in our model to two previously described bistable systems involving phages and bacteria. One example of alternative stable states in a phage-microbial ecosystem has been described in Ref. (21). Unlike in our model, where regime shifts change the composition of bacterial species, the ecosystem modeled in Ref. (21) switches between the states with and without phages. The main feature responsible for this switching behaviour is a decrease of adsorption coefficient of the bacterial host when nutrients become scarce. Similar to regime shifts in our ecosystem, the feedback between the nutrient concentration and the abundance of phages is at the core of this bistable behavior.

Perhaps the most celebrated example of a bistable system is the genetic switch operating inside a bacterial host of a temperate phage (22, 23). In a host of the prophage *λ* there is an intracellular competition between the dormant, lysogenic state dominated by the repressor protein C1 (24), and the virulent, lytic state dominated by the protein Cro (25). When Cro wins, it leads to production of a large number of phages, akin to the species *B*_2_ in our microbial ecosystem. High nutrient concentration in the environment typically favors the lytic state of the *λ*-host (26). Such lytic state is analogous to the *B*_2_-dominated regime in our ecosystem, also favored by high *C*. In this sense our ecosystem can be in the “dormant state” producing few phages when it is dominated by *B*_1_. When this state is exposed to a strong pulse of *P, C*, or *B*_2_ it can switch to the “lytic state” dominated by *B*_2_ and producing many phages (see Fig. 3).

One realistic implementation of bistability predicted by our model is in a phage-microbial ecosystem consisting of a bacterial strain protected against phages by the abortive infection (Abi) mechanism (*B*_1_) and a partially-resistant strain (*B*_2_) co-infected by the same phage. Hosts with abortive infection allow phages to enter and kill them without producing a noticeable phage progeny (13). An example of the Abi defense is provided by certain types of CRISPR defense (27, 28, 29), where phages kill most of infected hosts but have zero or small burst size. In contrast to Abi- or CRISPR-protected bacteria, partially resistant strains may arise due to a mutation in the receptor protein which reduces both the growth rate (30) and the phage adsorption but has little effect on the burst size. Thus regime shifts may naturally occur as a consequence of diverse phage defence mechanisms in microbial ecosystems (31).

A potential application of our system is in a new type of phage therapy in which phages targeting the pathogenic species (*B*_1_) are introduced together with carefully selected non-pathogenic species (*B*_2_) infected by the same phage. This therapy effectively combining two population pulses shown in panels A and C of Fig. 3 would lead to a more efficient and permanent elimination of the fast-growing pathogen (*B*_1_). One of the advantages of this approach is that phages would be continually present in the former patient thereby preventing reentry of pathogenic bacteria. The strategy could be made even more favourable if the bacteria added together with phages would use a nutrient other than *C* rendering it not vulnerable to nutrient competition from the pathogen.

## METHODS

### Simulations

The paper investigates the dynamics of a model defined by Eqs. (1-4), built on assumptions of mass action kinetics in a well-mixed system with an adjustable nutrient supply rate (32). We performed both deterministic and stochastic simulations of this model.

In stochastic simulations shown in Fig. 1B we use the Gillespie algorithm with step size of 0.0002 and rates defined for each of the 9 basic processes in Eqs 1-4: nutrient introduction and dilution events, *B*_1_ and *B*_2_ replication events, phage infection events separately in *B*_1_ and in *B*_2_, and combined death/decay/dilution events in each of the two bacteria and one phage species. Notice that a single phage infection event reduces the bacterial population by the step size equal to 0.0002, but increases the phage population by *β* · 0.0002. A large value of the burst size *β*_2_ = 40 justifies a small step size used in our simulations.

Deterministic simulations shown in Fig. 2 solve the dynamics given by Eqs. (1-4). At each value of nutrient supply rate *ϕ* we integrate the equations for 1000 time units to eliminate transients. We then increase the nutrient supply rate in increments Δ*ϕ* = 0.01. We use the steady state populations/concentrations obtained at *ϕ* as starting populations/concentrations for simulations at *ϕ* + Δ*ϕ*.

Each blue or red dot in Fig. 3 was obtained by starting the system in one of the stable states, and subsequently changing one of the variables (*P, C, B*_1_ or *B*_2_) as indicated on the y-axis. After a deterministic simulation of dynamical Eqs. (1-4) for 1000 time-units, the final state is compared to each of the states possible for a given value of *ϕ* and is marked with the corresponding color Fig. 3.

### Conditions for bistability

In our model it is convenient to describe the growth of a microbial species in (*C, P*) coordinates, characterizing respectively the nutrient and the phage concentrations in the environment. The population of a species exponentially grows for *λC* - *ηP* > *δ*_*B*_, exponentially decays for *λC* - *ηP* < *δ*_*B*_, and stays constant for *λC* - *ηP* = *δ*_*B*_. The last equation defines the so-called zero Net Growth Isocline (ZNGI) (14) of the species defined by all environmental parameters where the population of this species could be in a steady state. Everywhere in the region of the (*C, P*)-plane located to the right and below of species ZNGI (high *C* and small *P*) its population exponentially grows, while in the region to the left and above its ZNGI (low *C* and large *P*) it exponentially decays.

Red and blue straight lines in Fig. 4A correspond to the zero Net Growth Isoclines (ZNGI) of, correspondingly, the fast- and the slow-growing bacterial species in our model. They intersect at the point (*C* ^*^, *P* ^*^) given by

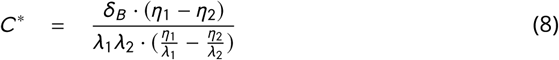

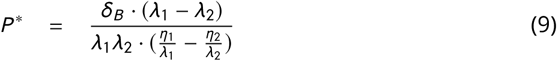

The intersection point correspond to the only set of environmental parameters at which these two species can potentially coexist with each other.

The lower part of the species 1 ZNGI (the solid part of the red line) extending from *P* = 0 and up to the intersection point at *P* ^*^ and the upper part of the species 2 ZNGI above *P* ^*^ (the solid part of the blue line) have a special property that the other species would not be able to grow in this environment. Hence, the union of these two halves of ZNGIs corresponds to uninvadable states of the ecosystem, which are the main focus of this study. The exact position of the environmental parameters on the (*C, P*) plane is determined by the supply rate *ϕ* of the limiting nutrient to the ecosystem. For *ϕ* < *δ*_*C*_ *δ*_*B*_ /*λ*_1_ there is not enough nutrient to support the growth of any species and the environment remains abiotic. Hence the first transition happens at

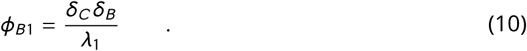

For *ϕ*_*B*1_ < *ϕ* < *δ*_*C*_ *δ*_*B*_ /*λ*_1_ + *δ*_*P*_ *δ*_*B*_ /(*Y*_1_*β*_1_*η*_1_), the species 1 is present but its biomass is not suffcient to support the survival of the phage. The phage first enters the ecosystem at

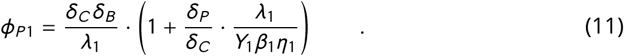

For even larger nutrient supply rates: 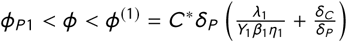 the ecosystem contains only the species 1 and the phage. The crucial parameters of the phage-bacterial ecosystem considered in our model are

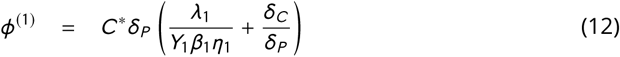

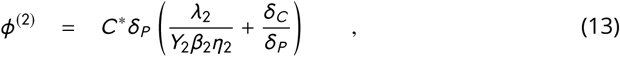

where *C* ^*^ is given by the Eq. 8. For nutrient supply rates *ϕ* > *ϕ*^(1)^ the species 2 can in principle grow in the ecosystem give *C* and *P* shaped by the species 1. What happens in this region crucially depends on whether *ϕ*^(1)^ < *ϕ*^(2)^ or *ϕ*^(1)^ > *ϕ*^(2)^, with the latter case corresponding to bistability which is the main focus of this study. For pedagogical reasons, let us first consider the model where *ϕ*^(1)^ < *ϕ*^(2)^ and thus *B*_1_-*B*_2_ co-existence is possible. In this case both species 1 and 2 can co-exist with each other in the interval 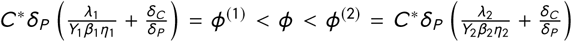. The abundances of each of the two microbial species can be geometrically determined as the intersection of two straight lines in the (*B*_1_, *B*_2_)-plane shown in Fig. 4B. The grey line corresponds to the steady state of the phage population *P* in Eq. 4 and is given by the equation

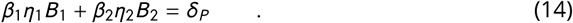

It must intersect with another straight line defining the steady state of the nutrient concentration *C* = *C* ^*^ and is given by

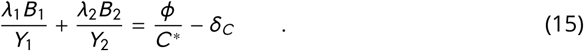

These lines intersect for positive *B*_1_ and *B*_2_ when *ϕ*^(1)^ < *ϕ* < *ϕ*^(2)^.

In the opposite case, where *ϕ*^(1)^ > *ϕ*^(2)^, the system is capable of bistability for nutrient supply rates *ϕ*^(2)^ < *ϕ* < *ϕ*^(1)^. To understand this it is useful to follow the trajectory of environmental parameters (*C, P*) as *ϕ* is gradually increased. For *ϕ*_*P* 1_ < *ϕ* < *ϕ*^(1)^ the environmental parameters follow the ZNGI of the fast growing species 1 (the red line in Fig. 4A below the intersection with the blue line). Immediately above the intersection point (*C* ^*^, *P* ^*^), realized for nutrient supply rate slightly larger than *ϕ*^(1)^, the ecosystem becomes invadable by the species 2. However, for this species the intersection point (*C* ^*^, *P* ^*^) corresponds to a lower value of nutrient supply *ϕ*^(2)^ < *ϕ*^(1)^. Hence after a brief transient period the environmental parameters (*C, P*) of our ecosystems move to the position marked with the blue cross in Fig. 4A. As *ϕ* continues to increase above *ϕ*^(1)^, the environmental parameters follow the ZNGI of the species 2 (the blue line to the right of the blue cross in Fig. 4A).

If at some point one starts decreasing *ϕ*, the species 2 will persist down to *ϕ*^(2)^ at which the environmental parameters are again at the coexistence point (*C* ^*^, *P* ^*^). For slightly lower *ϕ* the environmental parameters will discontinuously jump to the point marked with the red cross on the ZNGI of the species 1. For even lower nutrient supply rates they will continue to follow the ZNGI of the species 1 to the left and below of the red cross. Hence, our environment is bistable in the interval of two ZNGIs between the red and blue crosses. The lower red part of this interval is reachable only when *ϕ* is increased from a low value below *ϕ*^(2)^, while the upper blue part - when *ϕ* is decreased from a high value above *ϕ*^(1)^.

Above we assumed that phages can survive for *ϕ* = *ϕ*^(2)^ in the ecosystem dominated by the species 1 instead of species 2. This requires 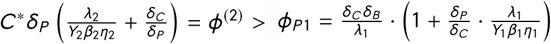, which can be rewritten as

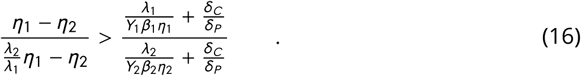

In the opposite limit of this inequality and for nutrient supply rates satisfying *ϕ*^(2)^ < *ϕ* < *ϕ*_*P*1_ the phages will be absent in one of the two alternative stable states (dominated by the species 1) but present in another one (dominated by the species 2).

The scenario illustrated in Fig. 2 corresponds to *ϕ*_*P*1_ < *ϕ*^(2)^ < *ϕ*^(1)^. In this case, the abundances in the steady state S dominated by the fast growing species 1 are given by

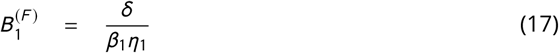

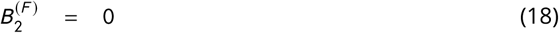

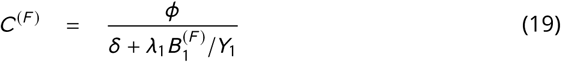

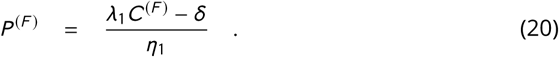

The abundances in the alternative stable state F dominated by the slow growing species 2

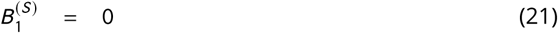

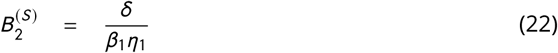

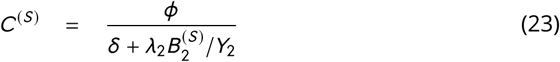

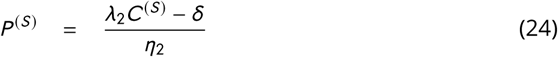

in the state dominated by the species 2.

In the regime where *ϕ*_*P*1_ < *ϕ*^(2)^ < *ϕ*^(1)^ and for nutrient supply rates in the bistable window *ϕ*^(2)^ < *ϕ* < *ϕ*^(1)^, the ecosystem also has a dynamically unstable steady state in which both bacterial species co-exists with each other and have the following abundances:

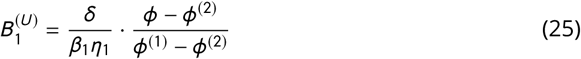

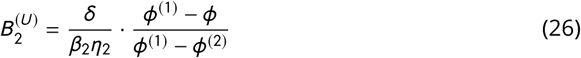

Note that in our study we consider only uninvadable states of the ecosystem. In other words, we ignore an invadable steady state, where for a small value of *ϕ* the ecosystem is populated only by the species 2, or another invadable steady state realized for a large value of *ϕ*, where the ecosystem has only the species 1. These states are located on invadable parts of each species’ ZNGI, which are to the right and below the ZNGI of the other species in Fig. 4A. Indeed, in these regions an arbitrary small inoculum of the invading species would exponentially grow and thereby disrupt the steady state of the ecosystem moving the environmental variables to a new point on the (*C, P*) plane.

### Parameters used in our numerical simulations

Both in stochastic and deterministic simulations of our model shown in Figures (1-3) we used the following parameters:

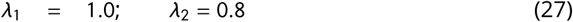

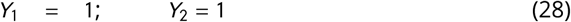

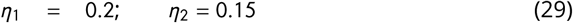

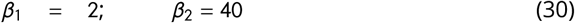

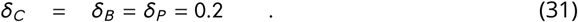

For these parameters the ecosystem is bistable when nutrient supply rate is between

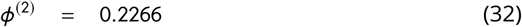

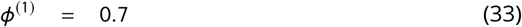

The bacterial abundances anywhere within this interval of nutrient supply rates are given by 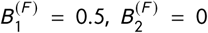 or 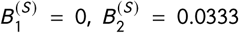 in alternative stable states dominated respectively by the fast and the slow-growing bacterial species.

The other transitions visible in Figure 2 happen at *ϕ*_*B*1_ = 0.04 above which the bacterial species 1 is able to survive given the dilution rate *δ*, and *ϕ*_*P*1_ = 0.14, above which the phage can survive in this ecosystem.

To estimate the typical values of *C* and *P* in two bistable states let us consider one example when *ϕ* = 0.25 is slightly above *ϕ*_(2)_. In this case the steady state concentrations of the nutrient and the phage in two alternative stable states: F and S are given by

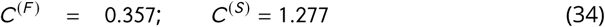

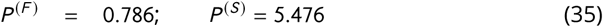

The dynamically unstable steady state point always has *C* ^*^ = 1 and *P* ^*^ = 4, which are located between their values in the F and S states. The bacterial abundances in an unstable state for *ϕ* = 0.25 are given by 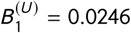 and 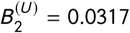. Note that the steady state abundance of the species 1 in the unstable state is much lower than its abundance 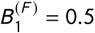 in the stable state. That suggests why for such a low value of *ϕ* we found it impossible to switch the ecosystem from the F state to the S state by pulses of *C, P*, or *B*_2_. Indeed, neither of these transient pulses is capable of lowering down *B*_1_ to the extra low saddle point value 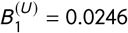 from initial stable state value of 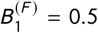 without simultaneously moving the populations of other species away from the saddle point region.

The position of the crosses in Fig. 4A can be calculated as follows: at *ϕ* = *ϕ*^(1)^ = 0.7 the species 1 sets the environmental parameters of the ecosystem exactly at the intersection point (*C* ^*^, *P* ^*^) = (1, 4) between ZNGIs of species 1 and 2. For slightly higher nutrient supply rates the species 2 eliminates the species 1 and the nutrient concentration shifts to 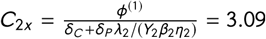, and the phage population - to *P*_2*x*_ = (*λ*_2_*C*_2*x*_ - *δ*_*B*_)/*η*_2_ = 15.14. On the way down the bacterial species 1 reenters the ecosystem slightly below *ϕ* = *ϕ*^(2)^ = 0.2266. When the species 1 replaces the species 2 immediately below this point the nutrient concentration shifts to 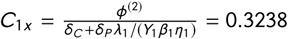, and the phage population - to *P*_1*x*_ = (*λ*_1_*C*_1*x*_ - *δ*_*B*_)/*η*_1_ = 0.6190.

## ACKNOWLEDGMENTS

This project has received funding from the European Research Council (ERC) under the European Union’s Horizon 2020 research and innovation programme under grant agreement No 740704.

